# Codistribution as an indicator of whole metacommunity response to environmental change

**DOI:** 10.1101/2022.06.24.497466

**Authors:** J. Christopher D. Terry, William Langdon, Axel G. Rossberg

## Abstract

Metacommunity structure can be summarised by fitting joint species distribution models and partitioning the variance explained into environmental, spatial and codistribution components. Here we identify how these components respond through time with directed environmental change and propose this as an indicator of sustained directional pressure. Through simulations, we identify how declines in the codistribution component can diagnose ecological breakdown, while rises in environmental and spatial components may indicate losses in peripheral areas and dispersal limitations. We test the method in two well-studied systems. Butterflies are known to be strongly responding to climate change, and we show that over 21 years the codistribution component declines for butterfly communities in southern England. By contrast, birds in the same region are under less climate pressure and, despite high occupancy turnover, show minimal change in metacommunity structure. The approach has high potential to summarise and compare the impact of external drivers on whole communities.

## Introduction

How can we tell when external environmental drivers, such as climate change, are impacting communities? Without further contextualisation, simply observing a change at a single location is insufficient, as communities are commonly subject to a considerable ongoing compositional turnover (Dornelas *et al*. 2013, 2014; Magurran *et al*. 2019) that is not necessarily associated with observable environmental change (Baselga *et al*. 2015). Community dynamism can be driven by ongoing environmental stochasticity as well as entirely endogenous processes (O’Sullivan *et al*. 2021) or ecological drift (Gilbert & Levine 2017). Separating ‘background’ community variability from that caused by directed environmental change is a considerable challenge, especially given that community reorganisation is not an instant process (Magurran *et al*. 2019). Metacommunity analyses, which examine multi-species responses across multiple sites, offer the potential to identify consistencies that could indicate an external driver.

Summarising high-dimensional (multi-species, multi-site, multi-time) data into informative metrics is a long standing, but crucial, challenge for understanding and addressing large scale ecosystem change. Clearly, answers to this question will depend on the type of data available. Where trait data related to environmental preference is available, community trait shifts can give clear indications of a directional driver (Wieczynski et al. 2019; Engelhardt et al. 2022). However, such data is not always available, and the overall picture that emerges may be complex as species within a community may respond differently (Antão *et al*. 2022). It is not always clear what the most relevant traits or environmental factors are, or if there even is a single key driving variable at all.

Recent developments in metacommunity ecology (Leibold & Chase 2018) provide a scaffold to increase the spatial and temporal scale of classic community ecology to address these problems. Much research effort has been directed towards identifying and partitioning the mechanisms that structure different metacommunities (Ovaskainen *et al*. 2019; Blanchard et al. 2020; Jabot *et al*. 2020). However, identifying even the relative contributions of different mechanisms from observations of metacommunities remains a considerable challenge and likely requires substantial temporal replication (Guzman *et al*. 2022). Joint Species Distribution Models (JSDMs) (Warton *et al*. 2015; Ovaskainen et al. 2016, 2017) are multi-variate regression models that provide a powerful tool to examine the complex patterns in spatially replicated community data. When JSDMs were initially proposed, the residual associations between species were identified as representing hypotheses for interactions between species. However, recent theoretical (Dormann *et al*. 2018; Zurell *et al*. 2018; Blanchet et al. 2020; Poggiato et al. 2021; Calcagno *et al*. 2022) and empirical (Barner et al. 2018; Freilich et al. 2018; Brazeau & Schamp 2019; Thurman et al. 2019) research has emphasized the poor correspondence with direct species interactions and the breadth of other factors that could determine the observed correlations in species responses. In particular, co-occurrences of species can be attributable not to direct species interactions, but rather to additional unmeasured environmental variables that are likely to obscure any underlying signal of species interactions (Blanchet *et al*. 2020).

Rather than seeing this as a shortcoming of JSDMs, we propose that this conflation of underlying mechanisms can be seen as an opportunity to detect the signal of unmeasured (or unmeasurable) environmental change on whole communities. Interspecific associations are increasingly recognised as an important component of biodiversity monitoring, and can be quantified in a very large number of ways (Keil et al. 2021). The JSDM-based metacommunity variance partitioning approach described by Leibold et al. (2022) summarises the structure of a metacommunity by partitioning community explanation three ways: into that provided by spatial association, known environmental determinants and codistribution between species. Here, we find in simulated mechanistic metacommunity models that directed environmental change is frequently characterised by a reduction in the explanatory power of the codistribution component, and increases in the spatial and environmental components. We then apply the method to two well-studied empirical systems. Like other JSDM tools, the overall approach is flexible enough to incorporate various sources of data, but importantly is capable of identifying signals of directed change without either trait data or direct knowledge of any putative driving variable.

## Method

The approach relies on fitting and comparing a series of JSDMs that predict the distribution of a community of species across a set of sites using environmental and spatial predictor variables. There are now numerous tools to fit JSDMs (see Wilkinson et al. (2019) for a review), although most differ principally in their approach to optimisation and, in principle, our approach is largely agnostic with respect to the optimisation approach used. Here we use species-by-site presence-absence data, since it is considerably more widely available, but we note that JSDMs can also fit abundance data, and this approach is likely applicable in that case too. However, we note that the statistical tools to assess the goodness of fit of count or abundance data are less well developed than for binary responses and further research may be necessary to identify possible unwanted biases in the variance partitioning. The method requires community data for a set of sites from at least two time points that are to be compared. We exclude species that fall below or above occupancy thresholds in any year to maintain sufficient variance in occupancy to represent an informative statistical target.

### JSDM fitting, variance partitioning and identifying shifts

We fit our JSDMs with a probit link function using the HMSC R package (v2.2) (Blanchet *et al*. 2019) that fits the models via MCMC optimisation. Latent variables identify residual codistribution of species as a separate element of the model (Warton *et al*. 2015; Ovaskainen *et al*. 2017) (Fig. 1a). We assume that the location of the sites is known and here represent spatial autocorrelation by the Moran Eigenvector Map (MEM) approach (Dray *et al*. 2012), decomposing the spatial coordinates using the *dbmem* function in the *adespatial* R package (Dray et al 2021) and selecting the first 10 components to use as spatial predictor variables. This approach has been found to be amongst the best at partitioning environmental and spatial drivers (Viana *et al*. 2022). More complex spatial structure, for example actual connectivity between sites, could in principle also be used but we do not discuss that further here. We assume that some environmental variables describing each site thought to be relevant to the distribution of the species in the metacommunity are known.

**Figure 1.**
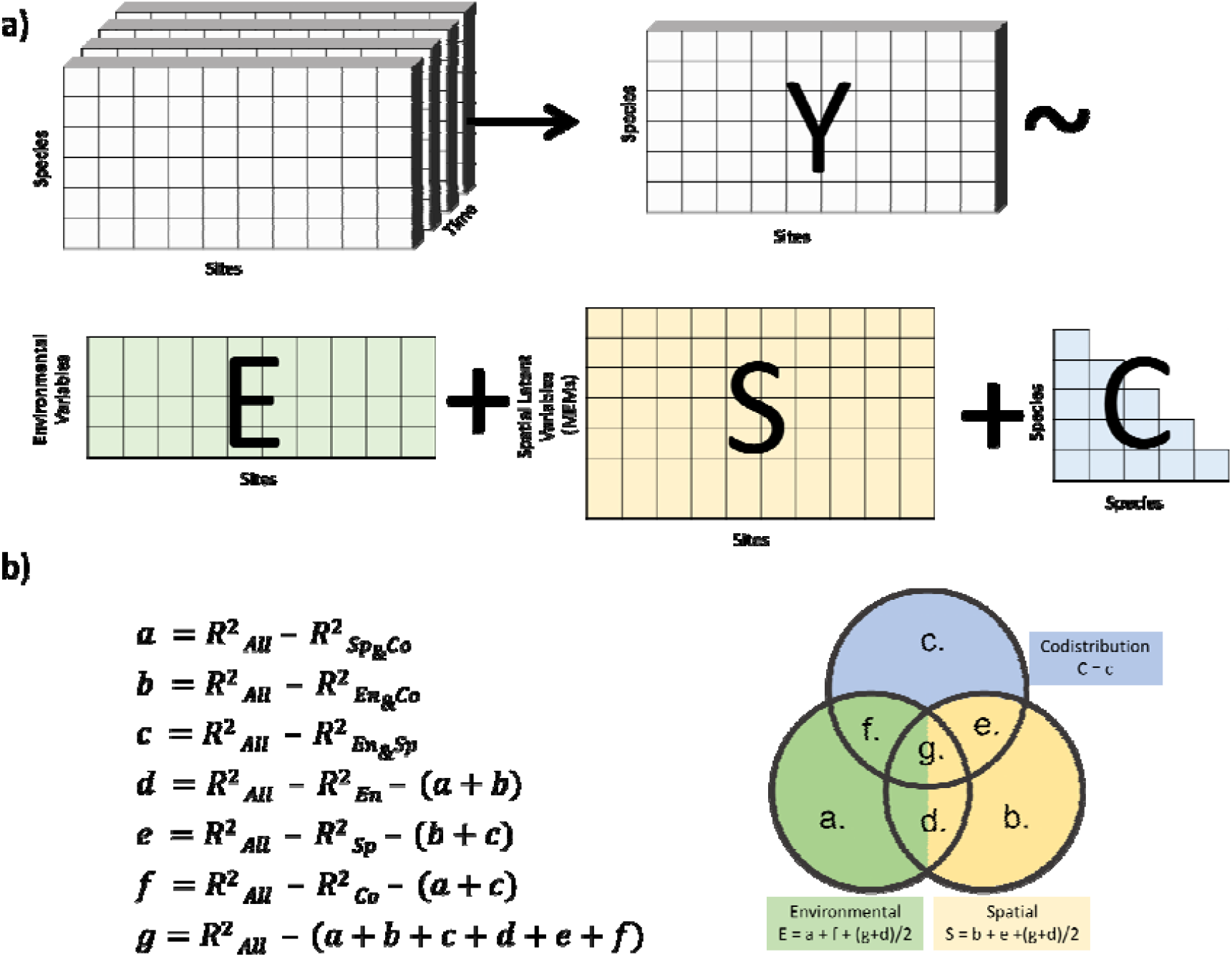
a). General illustration of inputs to our JSDM. For each time slice we obtain a *species x site* matrix (Y), that we wish to predict using some combination of environmental, spatial, and codistribution information. Note that codistribution between species is illustrated here by a correlation matrix between species, but, to reduce the number of parameters, we use a latent variable approach to summarise the correlation matrix into several latent factors for which each species has a factor loading (see Ovaskainen et al. (2017) for details). b) Having fit seven JSDMs using each possible combination of the three sets of information (*Sp, En, Co, Sp&En, Sp&Co, En&Co* and *All*), differences in the *R*^2^ of each model (calculated as per details in the main text following Leibold et al. (2022)) are used to partition the overall variance explained into sub-components (single effects and interaction terms, represented by sections of the Venn diagram). These are then summed into the three components of interest.

The variance partitioning is carried out following Leibold *et al*. (2022) by sequentially fitting seven models that make use of each possible combination of the spatial autocorrelation variables (Sp), measured environmental variables (En), and latent variables capturing species codistribution (Co), i.e. Sp, En, Co, Sp&En, Sp&Co, En&Co and ‘All’. For each model the total (i.e. including whichever random and fixed factors are included in the model in question) variance explained for each species is described using the conditional 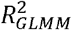 method (Nakagawa & Schielzeth 2013). Grouping some terms, this is calculated for each species separately as:

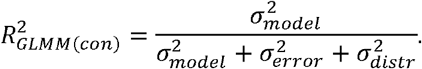

where 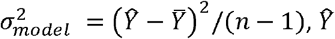 is the model prediction on the link scale (i.e. before transformation back from the unbounded scale to between 0 and 1), 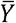 is the mean value on the linked scale, *n* is the number of sites, 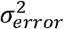 is the remaining variance unexplained by the model, and 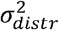 is the distribution specific variance - in the case of probit link functions this is 1 (Nakagawa & Schielzeth 2013). To make consideration for the way the models differ in the number of parameters they include, an adjusted-*R*^2^ metric is used that can handle latent variables (Gelman & Pardoe 2006):

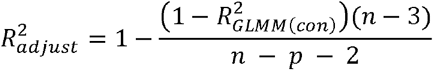

where *p* is the total number of fixed effects and latent variables estimated.

The fractional variance explained by each model component and interaction is determined using a Type III variance partitioning approach (Peres-Neto *et al*. 2006), working backwards from the most complex model to identify single effects and overlap terms (Fig. 1b). Then, to more efficiently summarise these quantities, we again follow Leibold *et al*.’s approach and group the sub-partitions of the total variance explained into three partitions of interest. Because latent variables are likely to efficiently capture residual variance the partitions overlapping with codistribution are assigned to the overall spatial and environmental partitions. Variance explained by the combination of environmental and spatial predictors is split 50:50 (Venn segments ‘g’ and ‘d’ in Figure 1b). To get a whole community value, we average across all the species (equally weighted). This variance partitioning process is repeated for each time slice, fitting each model independently. We then examine the changes through time in the variance explained by each of the three components. Simulation

To examine how the partitioning changes under known and controlled conditions, we used a mechanistic metacommunity model within a simple discrete-time Lotka-Volterra framework described in Box 1. These metacommunities are imagined to represent a set of sites within a wider landscape, and so are subject to both local dispersal within sites and some ongoing dispersal from ‘outside’ the observed system. We built a large set of distinct metacommunities, following a Latin-square design spanning 6 levels each of four parameters determining interspecific competition (*α*, 0.2:1.1), growth rates (γ, 5:50), dispersal (δ, 10^−5^:10^−2^), and environmental stochasticity (σ, 10^−3^:10^−1^), with two repeats at each combination (6^4^ × 2 = 2592 communities in total). Abundance data from just before the onset of the environmental change (year 975) and from after the onset of directional change (year 1000) from the central 100 sites of the arena was extracted and thresholded to presence/absence using a threshold biomass of 0.1. Species that had a total occupancy ≥5% and ≤95% in both time slices were retained (very rare or ubiquitous species have little occupancy variance that can be fit by the JSDMs).

#### BOX 1

**Mechanistic Metacommunity Simulator**

All communities were built on a square grid of 196 sites, each connected to their immediate neighbours (Fig. S1). Boundaries were connected as if on a sphere, so each site had four neighbours. The biomass of species *i* at site *z* at time *t* + 1 is given by:

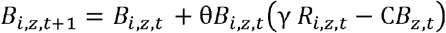

where *R*_*i*_,_*z*_,_*t*_ is the local growth rate, C is a competition matrix including inter and intraspecific competition, and θ is a timescale parameter, set to 0.1. γ is a simulation control parameter that controls the rate of intrinsic growth. The model is advanced in blocks of 10 t steps, which we refer to as a ‘year’.

Each site was assigned two explicit environmental variables (*E*_1_, *E*_2_). *E*_1_ represents a variable that will change through time, and could be considered ‘temperature’. It increases from left to right of the arena. Each year it is subject to additive stochastic noise drawn from a Gaussian distribution with mean 0 and standard deviation σ. *E*_2_ represents an environmental variable that stays constant through time, for example the underlying geology. It increases up the arena (Fig. S1). The growth rate of each species *R*_*i*_,_*z*_,_*t*_ is calculated using a cosine-based environmental performance function:

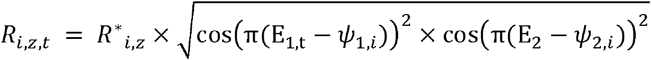

where *ψ*_1,*i*_ and *ψ*_2,*i*_ are species specific optima for each environmental variable. Additional species-specific site preferences are represented by also generating and incorporating a random species x site matrix *R*^*^. While complex, this function was chosen as it introduces moderate synergistic dependence on multiple environmental variables, it is always positive, its upper limit is controlled, and it approximately maintains the range size of each species within the full arena as *E*_1_ shifts.

Species are drawn from a global pool of 50 species each with randomly generated *ψ*_1,*i*_ and *ψ*_2,*i*_ values drawn from independent uniform distributions (0:1) and 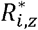 with uncorrelated entries drawn from a uniform distribution ranging from 0.2:1. The competition matrix C between these species is relatively sparse and semi-structured. Each species *i* exerts competitive pressure on the next species in the sequence (i.e. the lower off-diagonal). Additionally, random competitive effects are also introduced to C, with probability 0.01. All interspecific interactions have the same coefficient, α. The intraspecific (diagonal) elements are all set to 0.1.

Dispersal between sites and into the system is added at the start of every year. Local dispersal incoming into each site from the neighbouring sites *n* is calculated from *I*_*i*_,_*z*_,_*t*_ *δ*∑_*n*_*B*_*i*_,_*z*_,_*t*_. In addition, dispersal into the system from the global species pool is included. One species randomly selected from the global pool is each introduced to each site with probability 0.1 at density 0.0001.

Communities are initiated by starting a randomly selected half of the species in the global pool at a density of 1, each at a different randomly selected node. The simulation is run for a burn-in period of 975 years, during which the total species richness and range size distribution of the community reaches a steady state while the metacommunity continues to show ongoing turnover at local scales.

Directional environmental change is introduced by increasing *E*_1_ across the whole arena by 0.01 each year, for 25 model years. The corresponds to a displacement in the environmental envelope of each species equal to about half of the total range (see example in Fig. S1).

For each time slice, we fit separate JSDMs to conduct the variance partitioning as described above (total MCMC iterations 20’000, burn in = 10’000, thinning factor = 10, which trials suggested was more than sufficient for convergence). We fit the models using initial 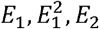 and 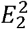 as environmental predictors. Observation error is known to be able to disrupt the accurate fitting of JSDMs (Beissinger et al. 2016). We examine the potential for this to be a problem in SI, by showing the overall robustness of the approach to the random introduction of false-negatives to the dataset.

A representative example (Fig. 2) shows the changes in metacommunity structure through time before and during the onset of climate change. Pre-climate change fluctuations in measured metacommunity structure are attributable to a mixture of background environmental stochasticity, intrinsic dynamics of the system (driven by competition and incoming dispersal), and a small amount of MCMC fitting error. With the onset of directed environmental change the community structure responds, after a brief delay, to reduce the codistribution contribution and increase the spatial contribution (i.e. moving to towards bottom right of the ternary plot). The system then settles around a new structure, as species continues to move through space in response to climate change.

**Figure 2.**
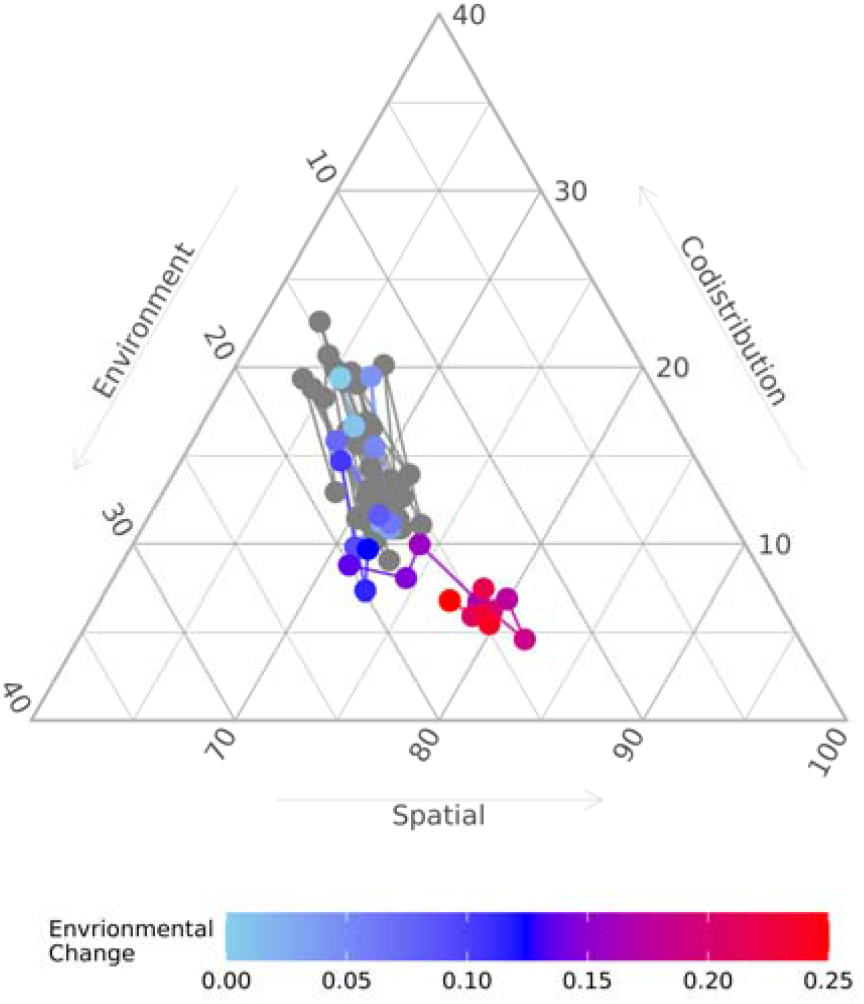
Metacommunity structure example simulated community through time before (grey) and during (coloured) the onset of directional environmental change. Each point on the ternary plot shows the percentage of the total *R*^2^ attributable to each partition. Grey points show the partition fit to 50 time points before the introduction of directed environmental change. Points are coloured when directional change in the variable *E*_1_ is introduced at a rate of 0.01 between each sample. Ternary plot made using ‘ggtern’ R package (Hamilton & Ferry 2018). Simulation model parameters used: *α* = 0.31, *γ* = 10.4, *δ* = 0.0028, *σ* = 0.01.

### UK Butterfly Transect Test

To examine if the pattern identified in the simulations is also discernible in real data, we used data from the United Kingdom Butterfly Monitoring Scheme (UKBMS). This highly curated data consists of 1-2 km long transects regularly surveyed by trained volunteers across the UK. Because UK lepidoptera are known to be undergoing climate-change driven range shifts, albeit influenced by landscape structure (Parmesan et al. 1999; Roy et al. 2001; Warren et al. 2001; Mair et al. 2012; Oliver et al. 2017), this dataset provides a ‘known target’. While the whole dataset covers surveys from 1976 to the present, we focus on data from 2000-2020 because in this period a large number of sites that have been continuously monitored.

We selected English transect sites which had been walked at least 5 times within at least 18 of the 21 years in our study period, and were south of 53.5° latitude and east of -3.5° longitude (thus excluding outlying peripheral sites in the west and north of England with distinct habitat types, Fig S6). We grouped individual transect walks within each year, to determine if each species was observed at least once in that year. We excluded moths (which are not consistently recorded at all sites), migratory species (*Colias croceus* and *Vanessa cardui*), and species with mean occupancies <2% or >98%. Species’ status during missing surveys (mean 1.57 years were missing per focal transect) were inferred based on whether each species was present in both the preceding and following surveys (inferred to be present in the gap), absent in both preceding and following (inferred to be absent). If the species was present at the site in just one of the preceding or following surveys, it was randomly assigned to be present with probability equal to its occurrence frequency at that site across the whole time period. This filtering process left 42 species and 97 sites. Overall occupancy through time was relatively constant (Fig. S7).

For each transect site, environmental information was gathered by cross-referencing the associated UK national grid cell (from the middle of the transect) with national databases. Habitat values of each site were inferred from the 2000 UK land cover map 1km raster (Fuller et al 2002). Although land cover changes are possible over the examined period, most sample sites are in relatively protected areas, and we do not expect widespread changes over the period. We generated four land cover variables for each site from summing key percentage land cover categories: Woodland (*‘Coniferous Woodland’*, +*’Broadleaved woodland’*), Farming (*‘Arable horticulture’* + *‘Arable cereals’* + *‘Improved grassland’* + *‘Arable non-rotational’*), Urban (*‘Continuous urban’* + *‘Suburban* / *rural developed’*), and *‘Calcareous Grassland’* on its own. The Woodland, Urban and Calcareous Grassland variables were log(x+1) transformed to better normalise their distribution. Calcareous grasslands are a key habitat type for European butterflies (van Swaay 2002), but within the CEH land cover map this is a broad habitat type that includes other high-pH grassland types. We therefore added a binary ‘chalk’ variable for each site based on the distribution of ‘*high*’ and ‘*high*(*variable*)’ in the carbonate content field of the British Geological Survey Soil Parent Material 1km dataset (Lawley 2012). As a baseline temperature variable, the mean average temperature of each site over the period 1961-1990 was inferred from the 5km HadUK-Grid (Hollis *et al*. 2019). The longitude and latitude coordinates of the selected sites were used to infer the first ten Moran eigenvector map components.

We conducted the tripartite variance partitioning separately for each of the 21 years in our sample period. For each submodel we ran very long chains (total MCMC sampling 100’000, burn-in = 50’000, thinning = 50). To confirm this was sufficient for convergence, we ran two MCMC chains of the full model and checked that all values of the Gelman-Rubin diagnostic 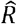 (Gelman & Rubin 1992) via the *coda* R package (Plummer *et al*. 2006). To assess the statistical significance of the trends, we fit separate linear models through each of the components.

### UK Breeding Bird Atlas Test

As a second test case, we used atlas data of UK breeding birds from the extensive 1970 and 2010 breeding bird surveys (Gillings et al. 2019). This is a highly curated citizen science dataset detailing the presence of all breeding birds in each 10×10km square of the UK national grid. Survey methods were calibrated to allow comparability between the 1970 and 2010 surveys. To exclude coastal-specialist species and focus on inland bird communities we used data from 348 grid cells from a rectangle of central England without coastline (51° : 54.3° latitude, -2.3° : -0.9° longitude, Fig. S11). We excluded non-native species (i.e not category A or C2 in the British list (McInerny et al. 2018)) and those that occurred in less than 5 focal cells, or were recorded as absent in less than 5 focal cells, in either year. This filtering process left 80 species. The total number of observed records in our dataset in each year was remarkably consistent (1970 = 12950, 2010 = 12798), but there was high turnover in occupancy patterns over this period, both within and between species (Gillings et al. 2019; Wayman *et al*. 2022). The breeding bird survey categorises the quality of observations into ‘confirmed’, ‘probable’ or ‘possible’. We repeated our analysis, excluding the ‘possible’ observations (which with the filtering procedure outlined above reduced the number of species to 75).

We determined environmental variables for each 10×10km grid cell from the 1990 (i.e. intermediate between the sample years) 1km grid CEH landcover map (Rowland 2020), calculating total percentage cover of 11 habitat variables using a mixture of specific target classes and some aggregate classes. These were: *Broadleaved woodland; Coniferous Woodland; Arable and Horticulture; Improved Grassland, Neutral Grassland; Calcareous Grassland; Acid grassland; Fen, Marsh and Swamp; Freshwater; Mountain, Heath and Bog; Urban and Suburban*. Elevation and climate variables correlated strongly with these habitats, and so were not included. The longitude and latitude coordinates of the selected sites were used to infer the first ten MEM components. Model fitting (total sampling: 50’000, burn in: 30’000, thinning: 20), convergence checking, and variance partitioning was conducted as for the UKBMS data, but with only two years we did not test for the significance of differences.

## Results

Across the full set of simulations, there was a clear tendency for the reduction in the codistribution component and a moderate trend for increases in the spatial and environmental components (Fig. 3, see also Fig. S4). The overall variance explained by the model showed minimal trend. Overall occupancy of the focal species x site matrix remained essentially constant with the change in environment (‘Before’: mean occupancy = 61.9%, ‘During’ mean occupancy = 62.2%), despite considerable turnover (Jaccard dissimilarity in each species’ site occupancy vectors between ‘before’ and ‘during’ samples mean = 0.306, SD = 0.099).

**Figure 3.**
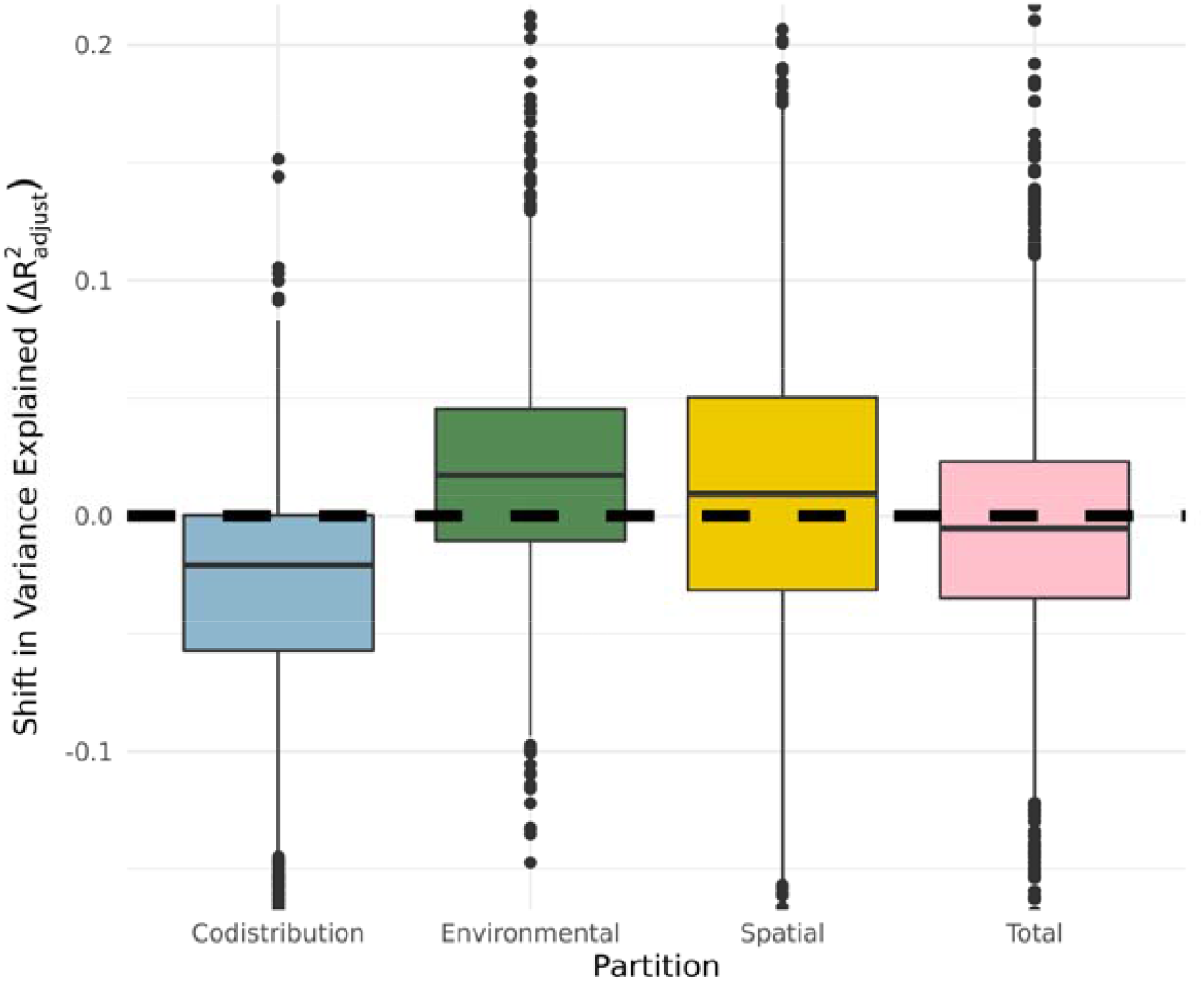
a) Distribution of changes in the partitions before and after the onset of climate change across the full set of 2592 simulated communities spanning a wide range of parameters. Boxplots hinges show 25 and 75^th^ percentiles, whiskers extend 1.5× inter-quartile range. Note that a few outliers have been clipped from the figure to maintain emphasis on the bulk of the distribution (largest observed shift was in codistribution, - 0.334).

The underlying dynamics (as determined by the four simulation control parameters) impacted the starting structure of the metacommunities (Figure S2 and S3). For example, higher levels of interspecific competition reduced the assembled diversity and average occupancy rates as well as increasing the variation in range size between species. Hence, higher competition also led to greater explanatory power for spatial structuring, while high levels of environmental stochasticity reduced the codistribution component.

However, simulation model parameters had relatively little influence on the shifts in structure induced by the environmental change (Fig. S4). The bulk of codistribution shifts were negative in all cases except for very high stochastic environmental variability where there was no consistent shift in any of the partitions. Shifts in the environmental partition tended to be positive across all parameter combinations. Shifts in the spatial partition were the most parameter sensitive, with the most consistent increased seen in parameterisations where the core growth rate was higher and competition was relatively weak. The introduction of false absences at high levels (e.g. 20%) led to a small reduction in the extent of the identified shifts in metacommunity structure (Fig. S5) but not the overall pattern.

Variance partitioning over each time slice in the butterfly dataset showed some support for a reduction in the overall codistribution partition (Fig. 4, -0.0013yr^−1^, p= 0.0575). Minor modifications in data filtering and predictor transformation can move the final p-value across the classic 0.05 critical threshold, but maintain the same trend. The environmental and spatial partitions showed no identifiable trend (Fig. 4). See supplementary information for species level variance partitions (Fig. S8), fitted environmental associations (Fig. S9) and species associations (Fig. S10).

**Figure 4.**
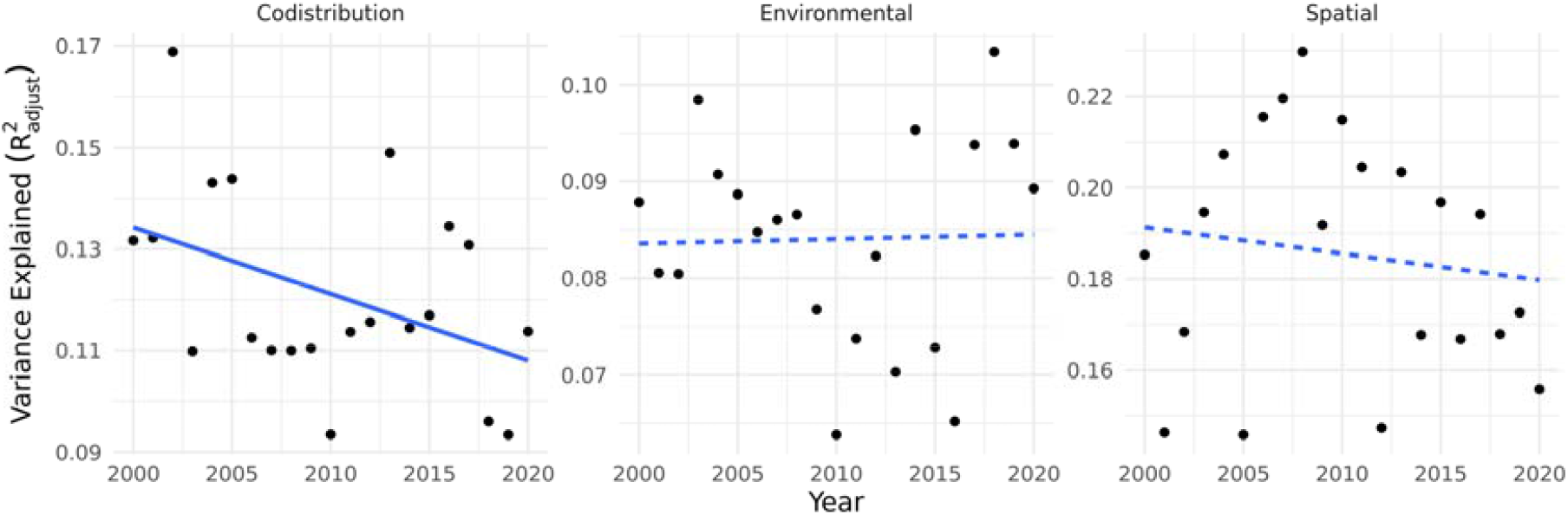
Trend in metacommunity structure of the UKBMS dataset through time. Lines show linear models. Codistribution shows some evidence for a negative trend (−0.0013yr^−1^, p= 0.0575), while other partitions showed minimal evidence of a linear trend (Environmental: p = 0.906, Spatial: p = 0.548).

The breeding bird dataset did not show large changes in overall metacommunity structure (Table 1), despite considerable occupancy turnover (27% of species x site occupancies changed between samples). Differences in codistribution structure were particularly small. Shifts in the environmental and spatial components were minimal and inconsistent depending on how the filtering was conducted.

**Table 1.** Comparison of the components of the variance partitioning of JSDMs fit to the breeding bird atlas datasets from central England between the 1970 and 2010 surveys.

| Dataset | Year | Total<br>Occupancy | Variance Explained |  |  |  |
| --- | --- | --- | --- | --- | --- | --- |
|  |  |  | Environmental | Spatial | Codistribution | Total |
| All records | 1970 | 46.5% | 0.333 | 0.131 | 0.088 | 0.552 |
|  | 2010 | 46.0% | 0.352 | 0.118 | 0.087 | 0.558 |
| Excluding<br>'possible' records | 1970 | 43.7% | 0.361 | 0.108 | 0.097 | 0.566 |
|  | 2010 | 40.3% | 0.330 | 0.110 | 0.095 | 0.536 |

## Discussion

Variance partitioning with JSDMs appears able to diagnose the impact of directed environmental change in complex communities. Despite continued challenges in the quest to identify underlying ecological processes from distribution snapshots (Gilbert & Bennett 2010), considerable progress is possible in other directions by seeking statistical features that indicate the impacts on community. As we will argue, the observed shifts have meaningful ecological explanations in line with the expectations associated with the early stages of community disintegration, the generation of no-analogue communities, and the role of dispersal limitations.

Our simulation model includes significant ongoing turnover, driven by continual immigration and competition, as well as underlying environmental stochasticity. This results in species occupancy frequency (and indeed the local species richness) remaining approximately constant as the environmental change is introduced, in line with observations from communities across the world (Dornelas *et al*. 2014). The observed shifts in the variance partitions can therefore be confidently identified as a signal of restructuring, not an artefact of loss.

Within the controlled virtual world of our models, the consistent pattern of metacommunity responses can have clear ecological explanations. The reduction in the explanatory power of species codistribution can be interpreted as being indicative of ongoing disruption to communities that reduces the consistency between species associations. As species distributions respond to the change in climate at different rates across sites, colonisation and extirpation stochasticity reduces the consistency of communities. There is some evidence that human action is introducing sustained changes to species associations through the Holocene (Lyons *et al*. 2016). We re-iterate that there is no reason to interpret this as a tendency for weakening effective interspecific interactions with climate change - there is little basis to identify changes in codistribution with any potential direct changes in pairwise biotic interactions induced by environmental change (Tylianakis *et al*. 2008; Poisot *et al*. 2015).

The frequent increase in the environmental partition can be identified as being driven by species being forced out of marginal habitats with environmental values on the edges of their original limits. Linked to this, the increase in spatial association is suggestive of dispersal limitation during the transient phase of ongoing environmental change. As isolated, peripheral, populations far from the species core areas are lost, newly colonised areas are likely to be closer to the core areas that provide the greatest source of new dispersers. The net effect is an increase in the explanatory power of spatial association as spatial heterogeneity is reduced. This pattern has been identified as a global response to current global change (McGill *et al*. 2015).

We would not expect real examples of communities subject to environmental change to precisely follow these same patterns: different guilds have different inherent traits, are structured differently by interspecific competition and are under different environmental pressures. Nonetheless, they can provide an expectation against which empirical communities can be assessed.

### Butterflies

The butterfly dataset showed some evidence for a decline in the codistribution partition, against a backdrop of year-to-year variation in abundance, indicating a weakening in the consistency of communities. Butterflies show large annual variation, responding to annual weather fluctuations (Roy *et al*. 2001). As such, the ability to identify a trend through a relatively short time series is a notable achievement of the methods. UK butterflies can be considered to not directly interact with each other – they rarely share larval food plants and possible apparent competition through shared parasitoids is not generally considered to be influential (Thomas *et al*. 2011; Thomas & Lewington 2014). The correlations between species can therefore be attributed with some confidence to unidentified variables. Inspection of the mean species association matrix (Fig. S10) suggests a split into a small group of woodland species from species more often associated with grassland. The habitat requirements of UK butterflies are well known (Redhead *et al*. 2016). However key factors, particularly the availability of specific larval food plants, are only loosely captured by our coarse environmental variables. The codistribution latent variables can capture correlated responses to unknown (or at least, unincluded) environmental variables. Given the expectation from our simulations that the environmental and codistribution components would respond in opposing directions, higher quality environmental predictors may be able to reveal stronger trends in both the residual codistribution and the environmental partitions. There was no hint of any change in the spatial association component through time. It is possible that dispersal limitation may not be relevant or detectable at the scale and resolution of our dataset. Butterfly dispersal capacity varies between species (Stevens et al. 2010), but in our dataset weaker dispersers, such as the Black Hairstreak *Satyrium pruni* (Thomas *et al*. 1992), are mostly excluded because of their rarity.

### Birds

UK birds are undergoing considerable change, but across the highly heterogenous community there would be less expectation that there are consistent driving variables compared to the butterfly dataset. To some extent, the bird data acts as a ‘negative control’ for our method. With just two time periods to compare, it is not possible to test statistically whether the changes in the spatial and environmental partitions represent a significant trend. However, we do find the apparent consistency, particularly in the codistribution component, worthy of note, especially within the context of the very high turnover rate. The fraction of observations listed only as only ‘*possible*’ confidence was higher in 2010, leading to a small decline in total occupancy in the stricter dataset that may have driven the reversal in the direction of change in spatial and environmental components. A reduction in the spatial partition (as we observed in the larger dataset) would be in line with the findings of Wayman *et al*. (2022) who found a moderate increase in spatial heterogeneity through time using similar data. However, if real, this may have different drivers to those examined in our simulations. As strong dispersers, birds are likely to be less subject to the dispersal constraints that drive the increase in spatial contribution in the simulations.

### Applicability and extensions

Given the ever-increasing number of systems to which JSDMs have been applied, we believe it is likely that this approach can be applied widely. A constraint we imposed is to fit each JSDM to the same sites and species through time, which limits the approach’s applicability to certain datasets where the coverage is large, but irregular. However, in principle this is not a necessity – if it can be reasonably assumed that the sampling of sites (and species) is consistent through time more heterogenous data sources could be used. Furthermore, while in both examples we used highly curated data sources where it is reasonable to assume that observation error is a small contributor, this may not be necessary. Although our simulations (Fig. S6) suggest that the approach is quite robust to false-absences, an important part of any deployment of the method will be testing for the imperfect detection of species (Tobler *et al*. 2019), which may be substantial and vary between both species and sites (Beissinger *et al*. 2016). Importantly, there is no particular need to have a pristine reference ‘before’ site with to which to compare. While longer time series are informative, shifts in communities are expected to continue and accelerate and ‘in progress’ shifts can also be detected.

That said, moving from summary statistics describing how a community is changing to diagnosing a causal link to specific external environmental drivers poses considerable challenges (Parmesan *et al*. 2013; Oliver & Morecroft 2014). Any measure of a metacommunity must always be considered in terms of the spatial scale at which it is observed (Leibold & Chase 2018), although this scale-dependence can be exploited to garner further ecological information (Ovaskainen *et al*. 2016; Elo et al. 2021; König *et al*. 2021). Examination of the responses of simulated models can clarify the expected direction of responses of metacommunities to different external pressures, and there remains significant scope for the exploration of different simulated models within this framework. Our measures of shifts in community structure are quantitative, but it is not yet clear how the magnitude of shifts in different systems (with different scales and environmental variables) can be compared. On the optimistic side, there are plenty of ecological explanations for these shifts – these are tangible quantities that are changing. On the pessimistic side, for the very reason that there are plentiful ecological explanations, identifying exactly which mechanisms are key is likely to require considerable additional information.

One way to mitigate the risk of generating ‘just-so’ stories derived from a limited number of summary variables is to examine the species level responses. While we have focussed on the whole-metacommunity responses as a general signal, this is constituted of the averaged responses of individual taxa which can also give useful information (i.e. the internal structure of the metacommunity sensu Leibold *et al*. (2022)). Investigation of the species level responses can bring additional insight into the drivers of the overall change. Here, we took the simplest possible route to weight the explanatory power within species within the community, taking a grand mean. Future work could profitably investigate if different weighting procedures can identify informative trends. While our focus here is on wider metacommunities and the possibility of inferring signals in systems with reduced data availability, an interesting avenue for future work would be to take advantage of the exceptional trait knowledge for both birds (Tobias *et al*. 2022) and butterflies (Middleton-Welling *et al*. 2020) to identify if drivers of species-level partitions can be identified.

### Conclusions

By providing a relatively simple summary of complex community level spatial-temporal trends, variance partitioning of JSDMs has great potential. Species associations identified through JSDMs have had a tumultuous recent history, with interpretations as interactions coming under particular considerable recent criticism (Blanchet *et al*. 2020). While they may not represent direct species interactions, they can still be highly informative at quantifying ecological responses. There remain very considerable challenges around interpreting ‘mechanistically’ what these ultimately statistical measures signify, but they have the potential to act as an early indicator of the road towards novel ecosystems (Hobbs *et al*. 2009) discernible through the noise of background turnover patterns.

## Acknowledgments

We thank the thousands of volunteers who contributed to the biodiversity datasets used in this work. Contains UK Butterfly Monitoring Scheme (UKBMS) data © copyright and database right Butterfly Conservation, the Centre for Ecology and Hydrology, British Trust for Ornithology, and the Joint Nature Conservation Committee, downloaded from the NBN Atlas. Contains British Geological Survey materials ©NERC Land cover, geology and climate data were accessed under an Open Government License

This research utilised Queen Mary’s Apocrita HPC facility, supported by QMUL Research-IT. http://doi.org/10.5281/zenodo.438045

JCDT and AGR are supported by the Natural Environment Research Council grant ‘Mechanisms and prediction of large-scale ecological responses to environmental change’ (NE/T003510/1). WL is supported by the Natural Environment Research Council Doctoral Training Partnership in Environmental Research (NE/S007474/1)

## Data availability

All code to reproduce the analyses and simulations, processed input data, and results summaries are available at https://github.com/jcdterry/MetacomPartition_Public. All raw empirical data used is open and available from the cited sources.

## Author Contributions

JCDT conceptualised the project, designed and ran the models, analysed the results, and wrote the manuscript. WL provided specialist advice on the empirical datasets. AGR obtained funding and contributed to the project design. All authors contributed substantially to manuscript editing.

