## Supplementary material for "Codistribution as an indicator of whole metacommunity response to environmental change": SI Figures

### SI Figures for *Codistribution as an indicator of whole metacommunity response to environmental change.*

Terry *et al.* (2022)

### Simulations

#### Simulation Setup Example

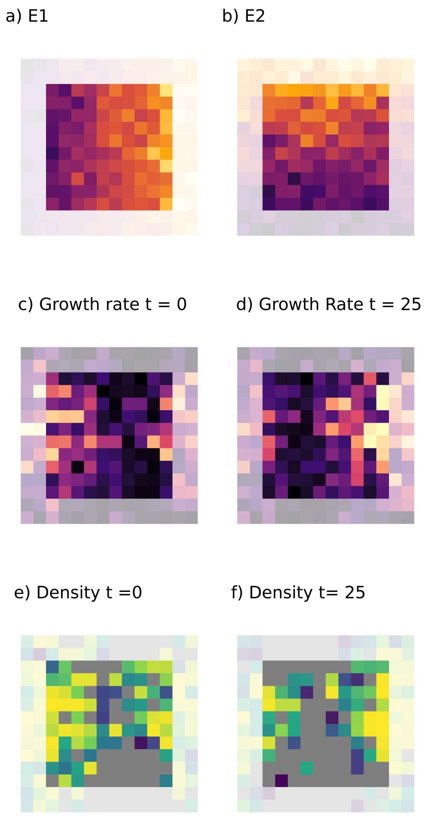

Figure 1: Example illustration of the simulation model grid and impact of directional change in the values of an environmental variable. Values are drawn from the same simulation as illustrated in the main text Figure 2. Values from the peripheral nodes not used for the JSDM fitting are shown greyed out. a) Initial distribution of $E_1$. b) Distribution of $E_2$. c) Growth rate $R$, of an example species before the onset of climate change. Note the approximately circular shape, but the high degree of heterogeneity. d) Growth rate R of the example species after 25 time steps of climate change ($t = 25$). Note the leftwards shift in the optimal (bright colours) habitat. e) Pre-climate change distribution of an example species. Note the approximate correspondence with the growth rates, but also infilling due to dispersal mass-effects f) Mid-climate change distribution of the example species. Note the movement lags - the shift leftwards movement is not as noticeable as in the growth rates (d).

#### Impact of parameters on metacommunity size and occupancy

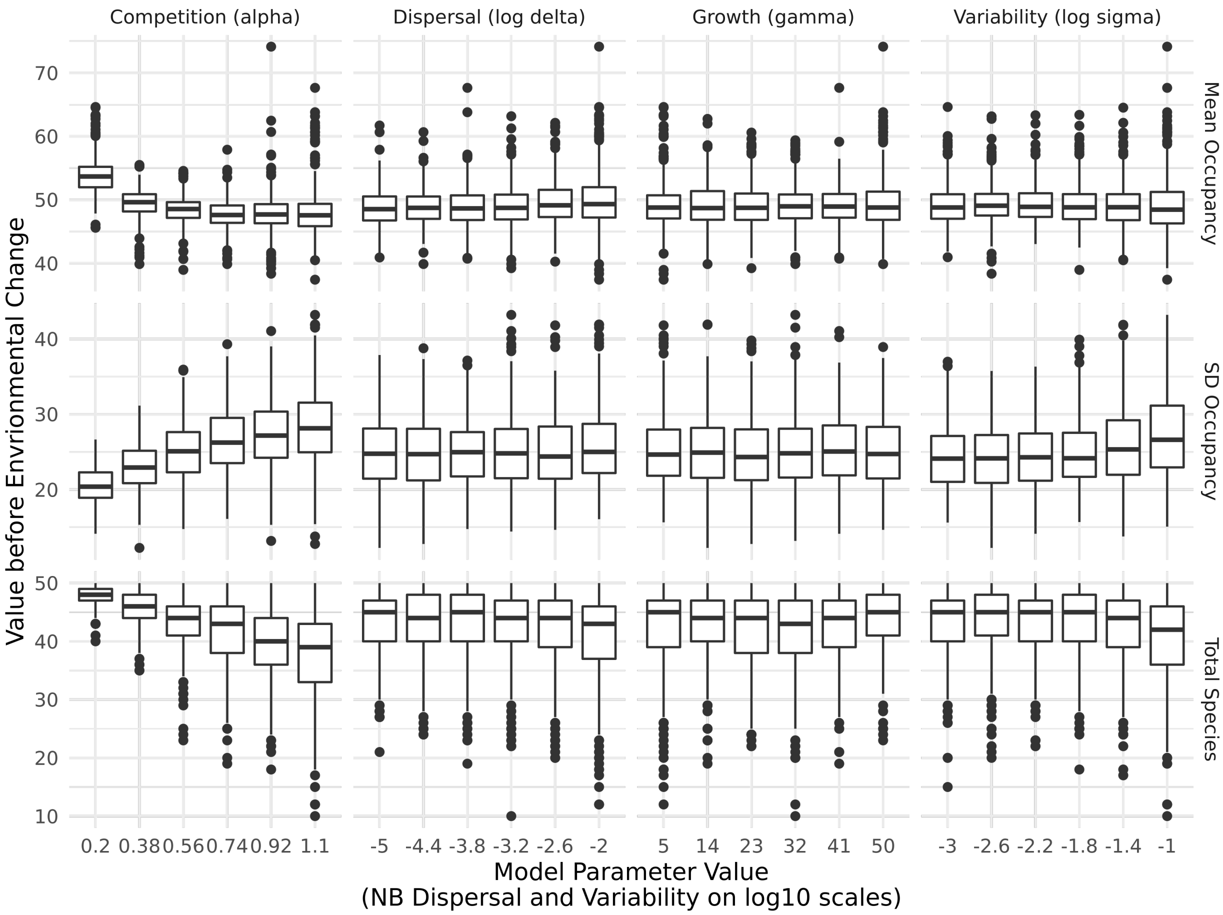

Figure 2: Impact of simulation model parameters on key metacommunity statistics prior to the introduction of environmental change. Only species that are used in the JSDMs are included here, i.e. excluding those that are too rare (or abundant) within the focal squares. Each facet includes all 2460 simulations, but are separated by different responses and driving parameters. Boxplots hinges show 25 and 75th percentiles.

#### Impact of parameters on simulated metacommunity structure

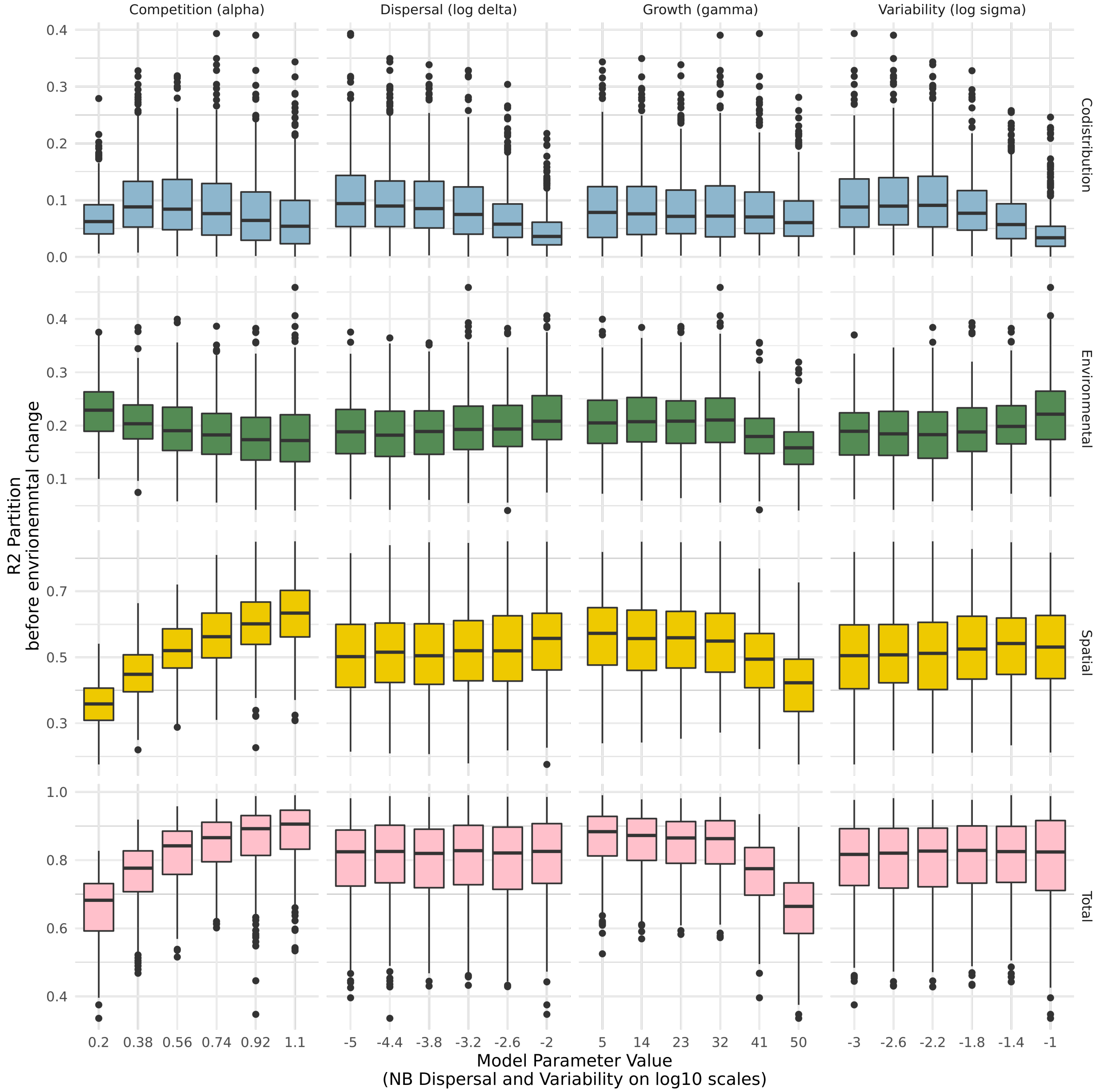

Figure 3: Impact of the model parameters on the simulated metacommunity structure, as assessed by variance partitioning before the introduction of environmental change. Boxplots hinges show 25 and 75th percentiles.

#### Impact of parameters on shift in simulated metacommunity structure

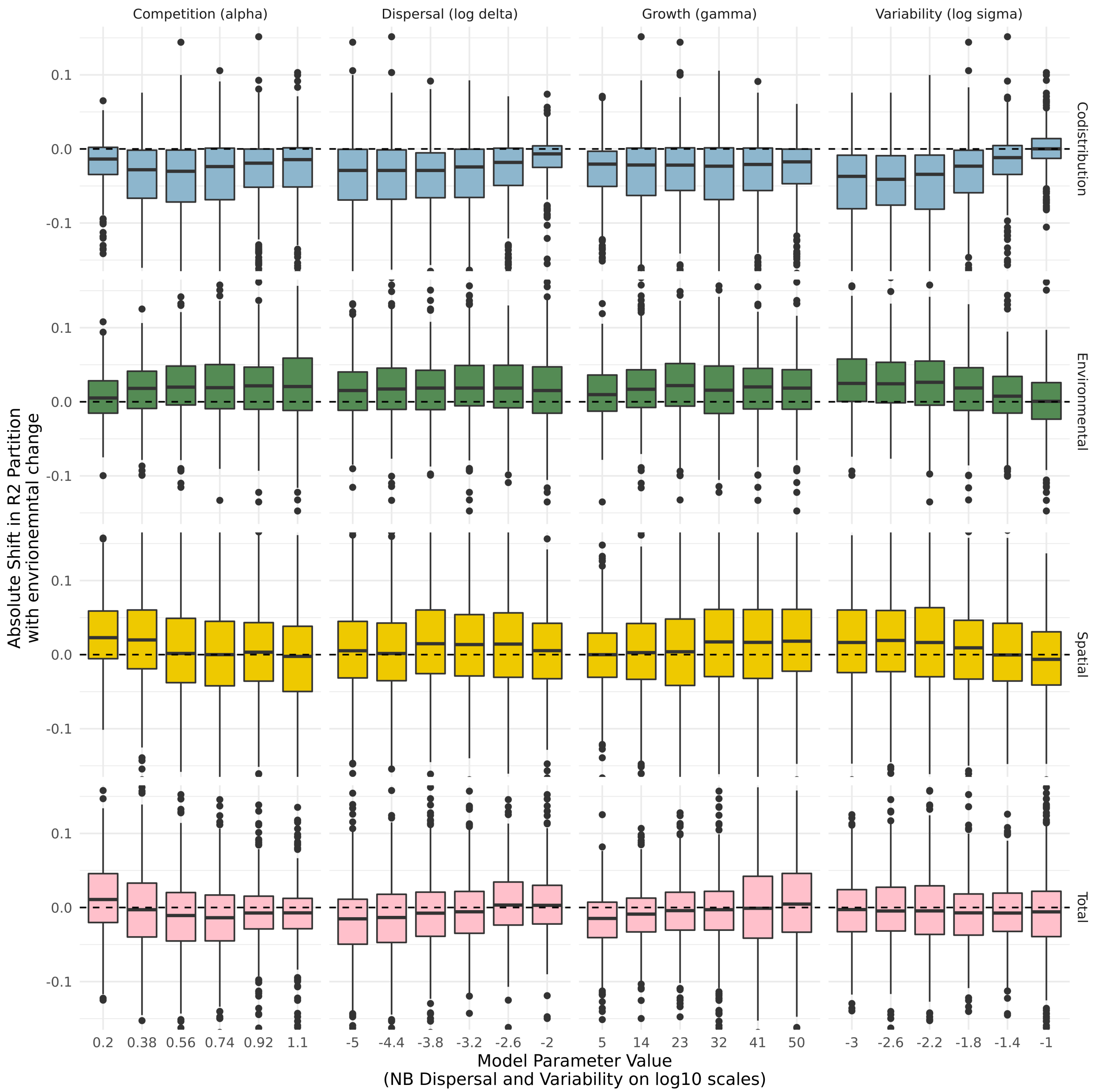

Figure 4: The dependence of shifts in the variance partitioning on the parameters of the simulated metacommunities. Figure 3 in the main text is a summary of this plot. Boxplots hinges show 25 and 75th percentiles.

**Impact of false absences on detectability of shift in metacommunity structure**

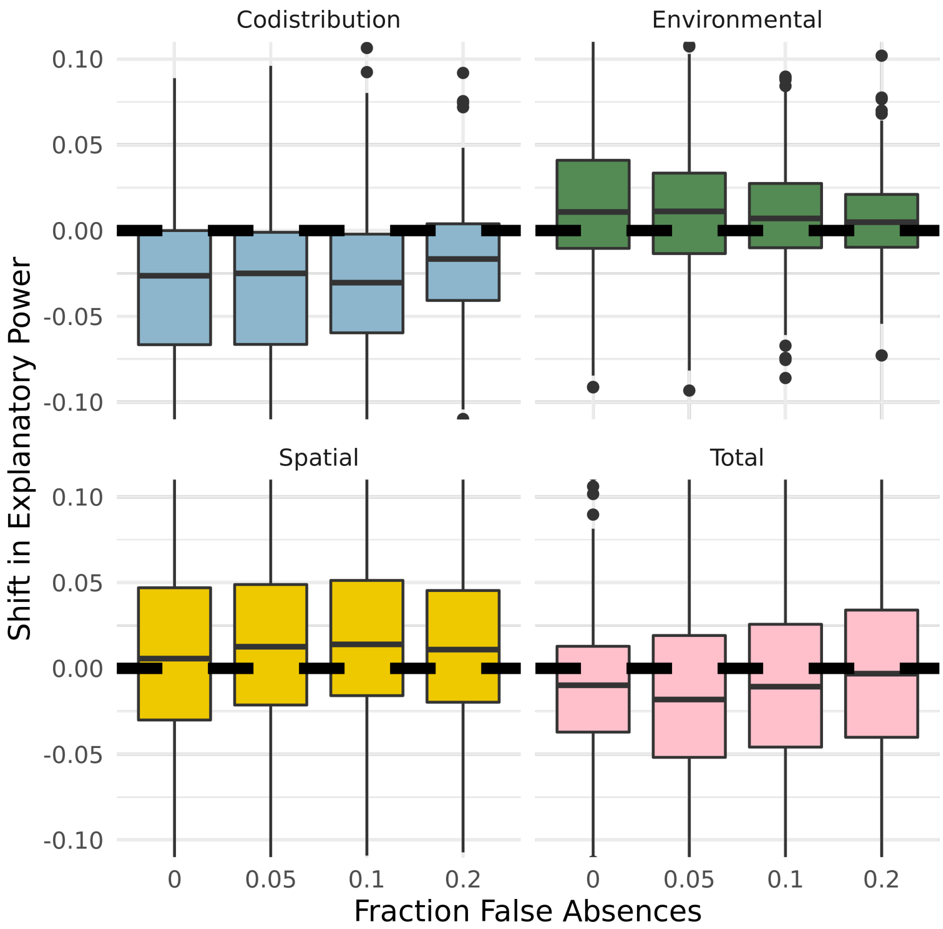

Figure 5: Impact of false absences on the consistent identity of trends. Simulation was run as with the analysis in the main text, but with a reduced spread of core model parameters ($\delta$: $10^{-5}, 10^{-4}, 10^{-3}$; $\alpha$; $0.3, 0.8, 1.1$, $\gamma$: $5, 20, 40$, $\sigma$: $10^{-3}, 10^{-2}, 10^{-1}$), crossed with 4 levels of false absences ($0, 0.05, 0.1, 0.2$). False absences are introduced by randomly, and independently, switching each presence (i.e. above threshold) to an absence with a given probability. Boxplots hinges show 25 and 75th percentiles.

### Butterfly Dataset

#### Distribution of Sites

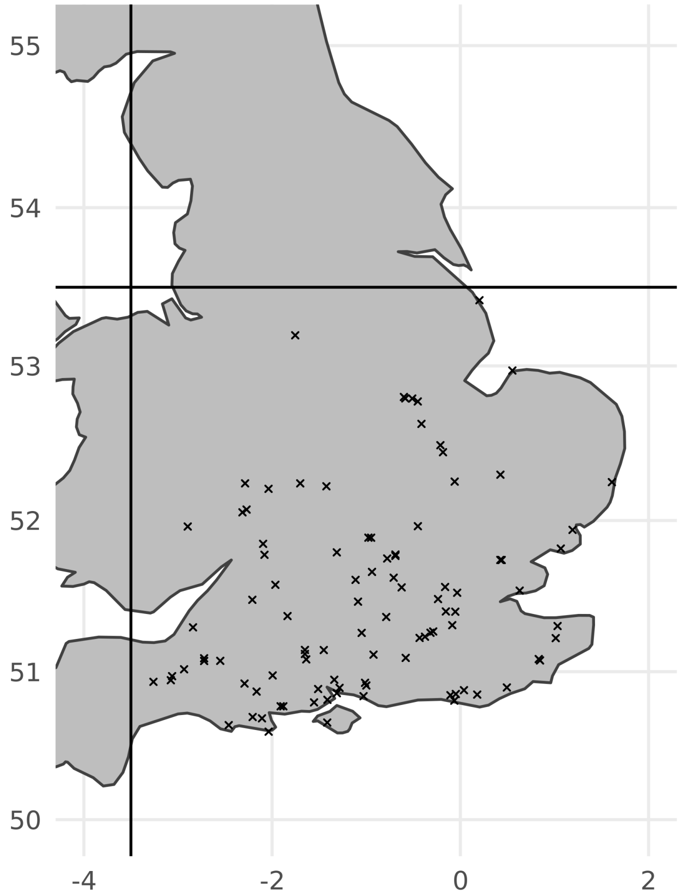

Figure 6: Distribution of UKBMS transect sites with sufficient data over focal period to be included in the analysis.

##### Occupancy Through Time

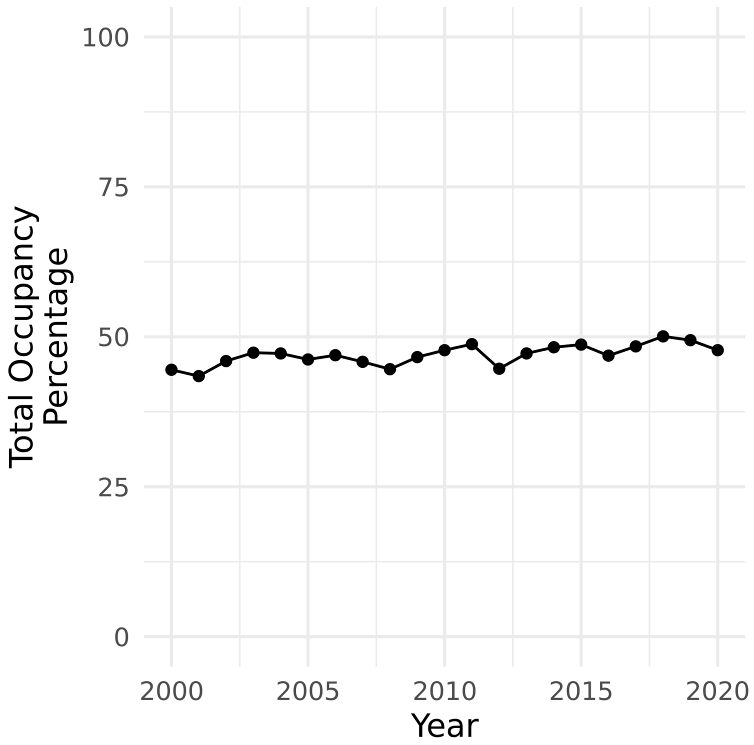

Figure 7: Total occupancy (i.e. total species:site presence records out of maximum possible) of focal butterfly species in focal sites across the time period.

#### Species-Level Responses

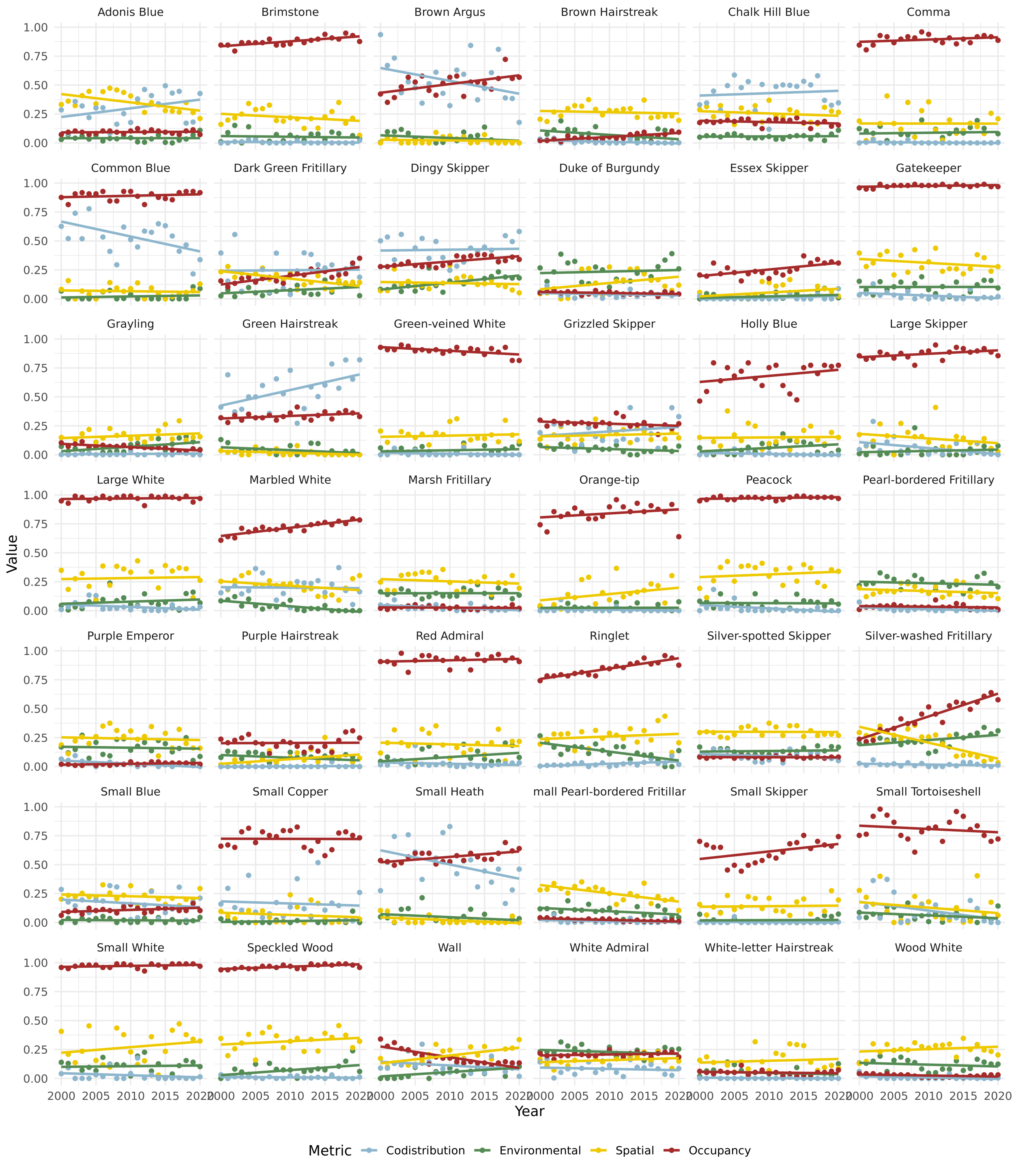

Figure 8: Species-level variance partitioning and species occupancy fraction (out of the 97 focal sites) through time. Note that the changes in the overall variance partitioning are driven by changes in relatively few of the species. However, these species (for example Common Blue and Green Hairstreak) are not necessarily those that are notably changing occupancy through this period.

#### Species Names

Table 1: Linnean binomials and English common names of focal butterfly species

| Scientific Name | English Common Name |
|---|---|
| *Polyommatus bellargus* | Adonis Blue |
| *Gonepteryx rhamni* | Brimstone |
| *Aricia agestis* | Brown Argus |
| *Thecla betulae* | Brown Hairstreak |
| *Polyommatus coridon* | Chalk Hill Blue |
| *Polygonia c-album* | Comma |
| *Polyommatus icarus* | Common Blue |
| *Speyeria aglaja* | Dark Green Fritillary |
| *Erynnis tages* | Dingy Skipper |
| *Hamearis lucina* | Duke of Burgundy |
| *Thymelicus lineola* | Essex Skipper |
| *Pyronia tithonus* | Gatekeeper |
| *Hipparchia semele* | Grayling |
| *Pieris napi* | Green-veined White |
| *Callophrys rubi* | Green Hairstreak |
| *Pyrgus malvae* | Grizzled Skipper |
| *Celastrina argiolus* | Holly Blue |
| *Ochlodes sylvanus* | Large Skipper |
| *Pieris brassicae* | Large White |
| *Melanargia galathea* | Marbled White |
| *Euphydryas aurinia* | Marsh Fritillary |
| *Anthocharis cardamines* | Orange-tip |
| *Aglais io* | Peacock |
| *Boloria euphrosyne* | Pearl-bordered Fritillary |
| *Apatura iris* | Purple Emperor |
| *Favonius quercus* | Purple Hairstreak |
| *Vanessa atalanta* | Red Admiral |
| *Aphantopus hyperantus* | Ringlet |
| *Hesperia comma* | Silver-spotted Skipper |
| *Argynnis paphia* | Silver-washed Fritillary |
| *Cupido minimus* | Small Blue |
| *Lycaena phlaeas* | Small Copper |
| *Coenonympha pamphilus* | Small Heath |
| *Boloria selene* | Small Pearl-bordered Fritillary |
| *Thymelicus sylvestris* | Small Skipper |
| *Aglais urticae* | Small Tortoiseshell |
| *Pieris rapae* | Small White |
| *Pararge aegeria* | Speckled Wood |
| *Lasiommata megera* | Wall |
| *Satyrium w-album* | White-letter Hairstreak |
| *Limenitis camilla* | White Admiral |
| *Leptidea sinapis* | Wood White |

#### Fitted Environmental Coefficients

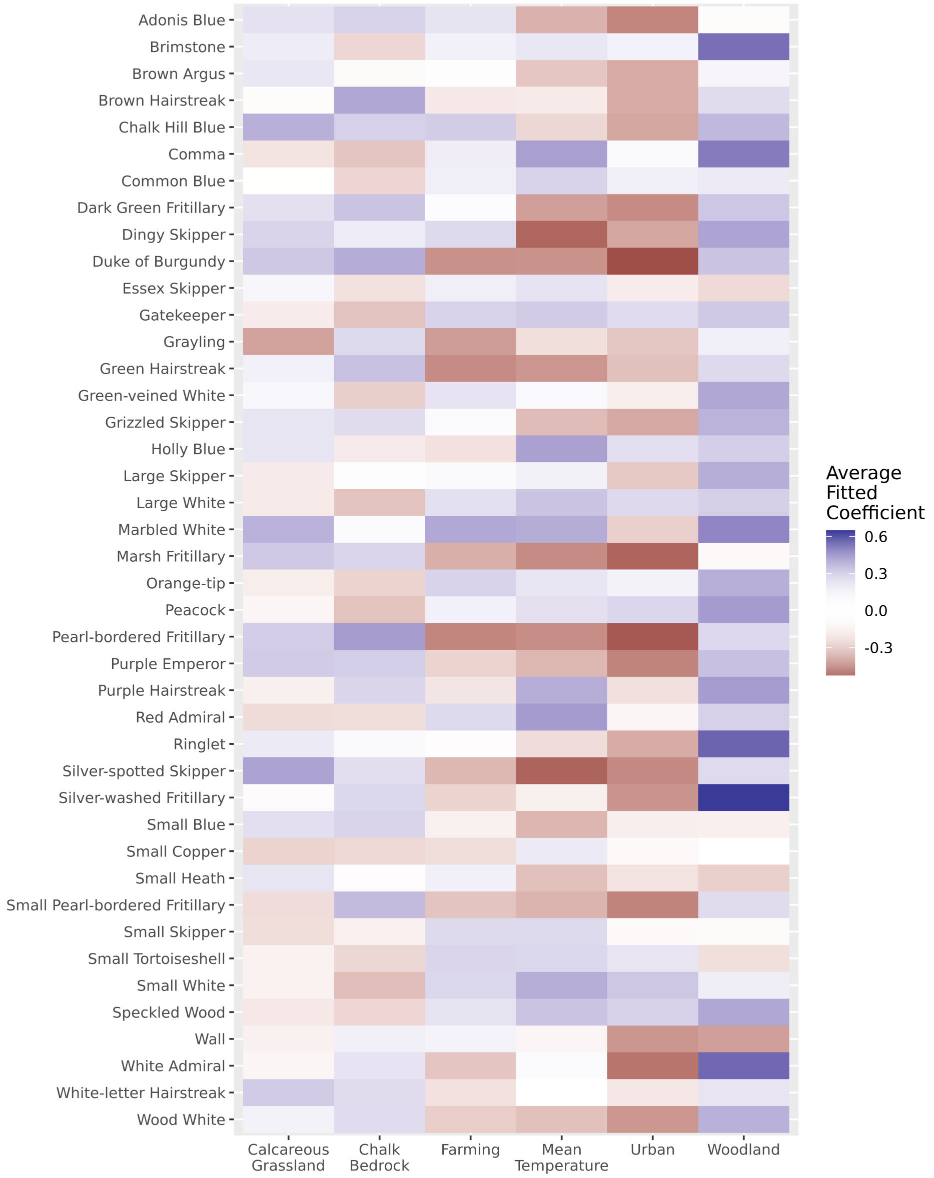

Figure 9: Butterfly fitted species-level environmental coefficients, averaged across all 21 years.

#### Fitted Species Associations

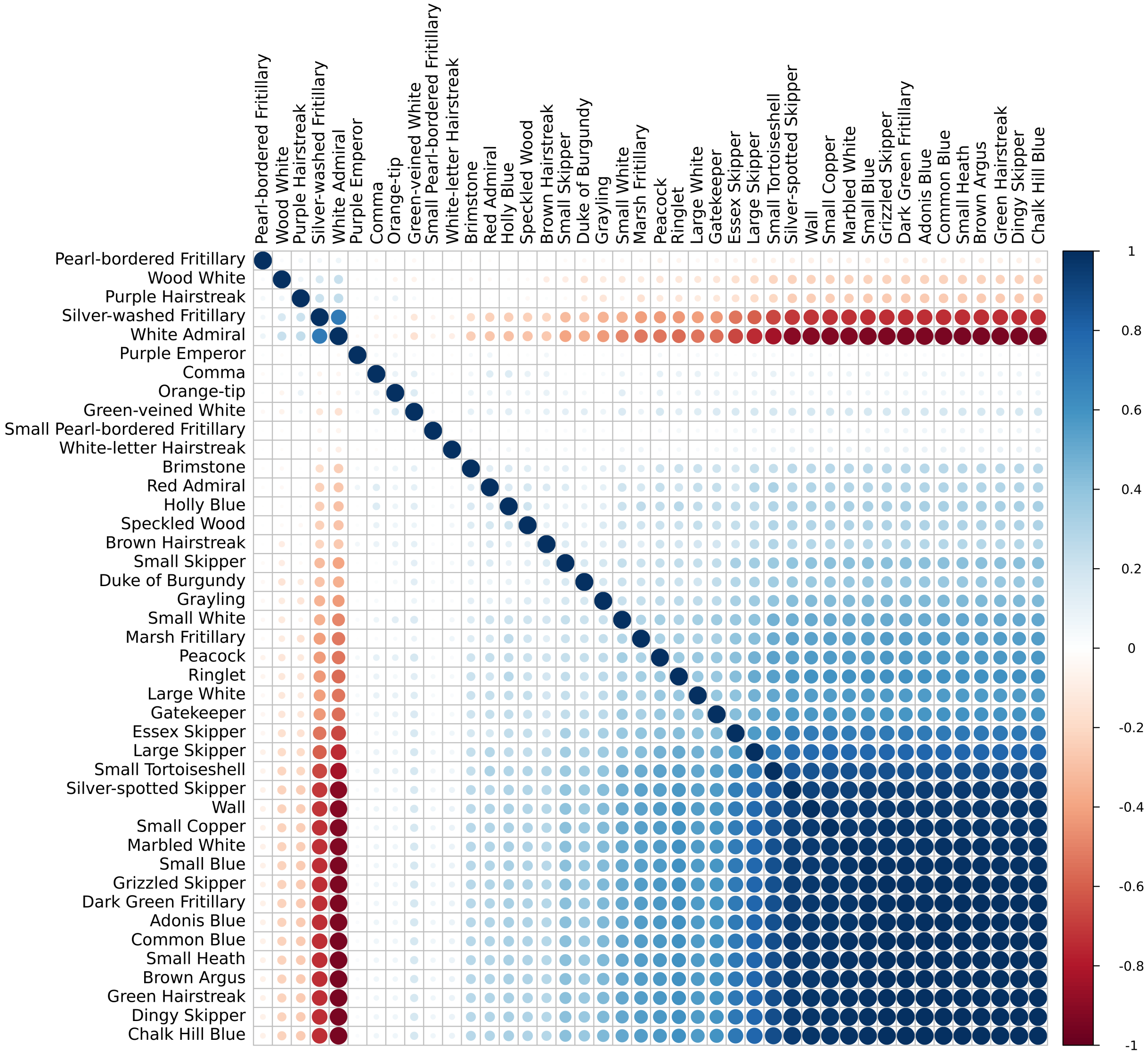

Figure 10: Fitted residual species associations defined by a correlation matrix, averaged across all time slices. Note the grouping into a large cluster dominated by chalk grassland species and a smaller cluster of species associated with woodlands.

#### Confirming Model Convergence

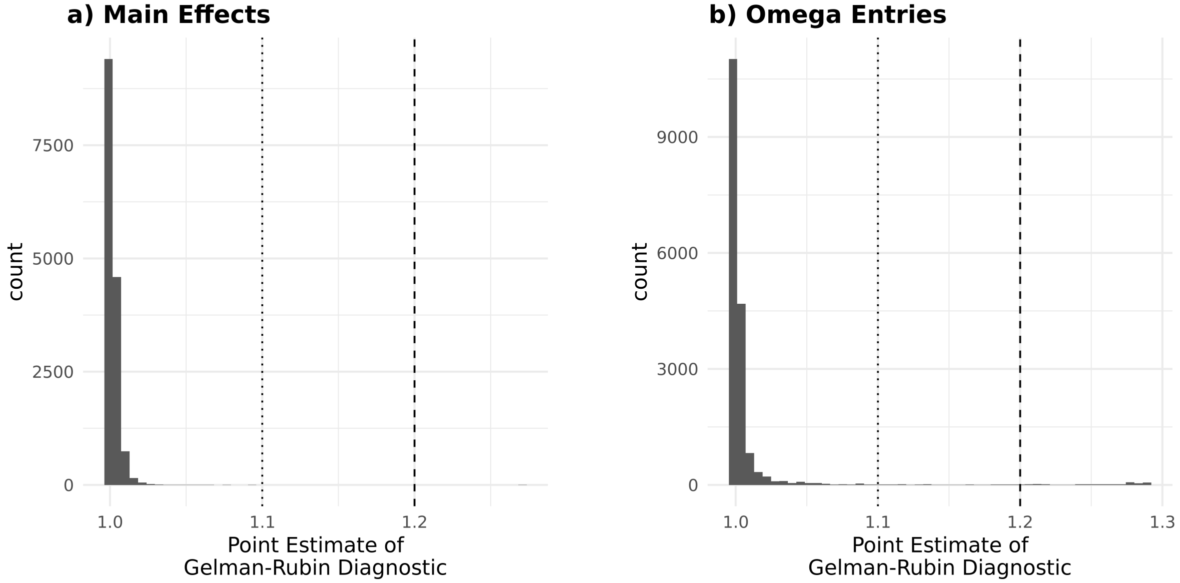

Figure 11: Histogram of Gelman-Rubin MCMC convergence statistics across all years of the butterfly dataset based on two independent chains fit for each year of the full model (total iterations = 100000, burn-in = 50000, and thinning = 50) a) Main effect coefficients (i.e. environmental and spatial coefficients) are all well below the standard threshold of 1.1, indicating acceptable convergence. b) Equivalent results for the species codistribution fitting are by necessity slightly more derived, as they are fit by latent variables that might not necessarily be fit in the same order, even if they converge. We therefore examine the convergence in the elements of the correlation matrix $\Omega$. Here the vast majority are well converged, although there are a few correlations that exceeded 1.2. However, as these were very much a minority and were not signifcantly over (the maximum was 1.29), we considered these models suitable converged.

#### Birds

##### Distribution of Sites

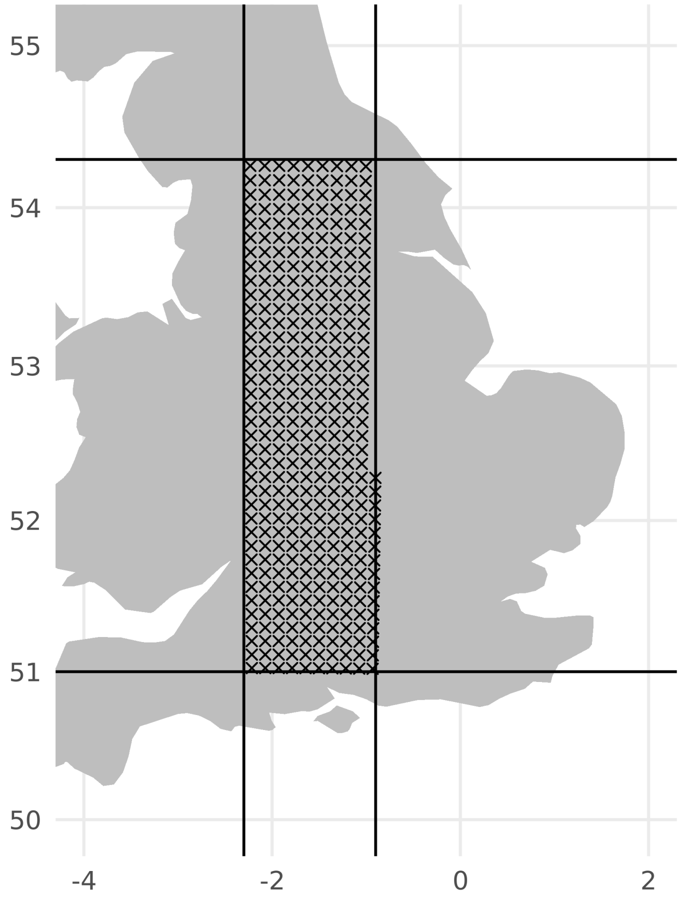

Figure 12: Location of the hectads retained from the British Breeding Bird Atlas dataset. Boundaries (51° : 54.3° latitude, -2.3° : -0.9° longitude) were chosen to be a simple shape that excludes coastal regions. The slight shoulder is due to the non-exact alignment between the UK National Grid and lines of longitude.

#### Species Names

Table 2: Linnean binomials and English common names of focal bird species. "HQ" column indicates if the species had sufficient "high-quality" observations to be retained in the more restricted datasets.

| Scientific Name | English Common Name | HQ |
|---|---|---|
| *Tyto alba* | Barn Owl | TRUE |
| *Chroicocephalus ridibundus* | Black-headed Gull | TRUE |
| *Lyrurus tetrix* | Black Grouse | FALSE |
| *Phoenicurus ochruros* | Black Redstart | TRUE |
| *Branta canadensis* | Canada Goose | TRUE |
| *Actitis hypoleucos* | Common Sandpiper | TRUE |
| *Sterna hirundo* | Common Tern | TRUE |
| *Fulica atra* | Coot | TRUE |
| *Emberiza calandra* | Corn Bunting | TRUE |
| *Numenius arquata* | Curlew | TRUE |
| *Cinclus cinclus* | Dipper | TRUE |
| *Calidris alpina* | Dunlin | TRUE |
| *Mareca strepera* | Gadwall | FALSE |
| *Spatula querquedula* | Garganey | TRUE |
| *Regulus regulus* | Goldcrest | TRUE |
| *Pluvialis apricaria* | Golden Plover | TRUE |
| *Accipiter gentilis* | Goshawk | TRUE |
| *Locustella naevia* | Grasshopper Warbler | TRUE |
| *Podiceps cristatus* | Great Crested Grebe | TRUE |
| *Picus viridis* | Green Woodpecker | TRUE |
| *Ardea cinerea* | Grey Heron | TRUE |
| *Perdix perdix* | Grey Partridge | TRUE |
| *Motacilla cinerea* | Grey Wagtail | TRUE |
| *Anser anser* | Greylag Goose | TRUE |
| *Coccothraustes coccothraustes* | Hawfinch | TRUE |
| *Circus cyaneus* | Hen Harrier | FALSE |
| *Falco subbuteo* | Hobby | TRUE |
| *Garrulus glandarius* | Jay | TRUE |
| *Alcedo atthis* | Kingfisher | TRUE |
| *Larus fuscus* | Lesser Black-backed Gull | FALSE |
| *Acanthis cabaret* | Lesser Redpoll | TRUE |
| *Dryobates minor* | Lesser Spotted Woodpecker | TRUE |
| *Sylvia curruca* | Lesser Whitethroat | TRUE |
| *Tachybaptus ruficollis* | Little Grebe | TRUE |
| *Charadrius dubius* | Little Ringed Plover | TRUE |
| *Asio otus* | Long-eared Owl | TRUE |
| *Poecile palustris* | Marsh Tit | TRUE |
| *Anthus pratensis* | Meadow Pipit | TRUE |
| *Falco columbarius* | Merlin | TRUE |
| *Cygnus olor* | Mute Swan | TRUE |
| *Luscinia megarhynchos* | Nightingale | TRUE |
| *Caprimulgus europaeus* | Nightjar | TRUE |
| *Sitta europaea* | Nuthatch | TRUE |
| *Haematopus ostralegus* | Oystercatcher | TRUE |
| *Ficedula hypoleuca* | Pied Flycatcher | TRUE |

| Scientific Name | English Common Name | HQ |
|---|---|---|
| *Aythya ferina* | Pochard | TRUE |
| *Coturnix coturnix* | Quail | TRUE |
| *Corvus corax* | Raven | FALSE |
| *Lagopus lagopus* | Red Grouse | TRUE |
| *Tringa totanus* | Redshank | TRUE |
| *Phoenicurus phoenicurus* | Redstart | TRUE |
| *Emberiza schoeniclus* | Reed Bunting | TRUE |
| *Acrocephalus scirpaceus* | Reed Warbler | TRUE |
| *Turdus torquatus* | Ring Ouzel | TRUE |
| *Charadrius hiaticula* | Ringed Plover | TRUE |
| *Columba livia* | Rock Dove | TRUE |
| *Riparia riparia* | Sand Martin | TRUE |
| *Acrocephalus schoenobaenus* | Sedge Warbler | TRUE |
| *Tadorna tadorna* | Shelduck | FALSE |
| *Asio flammeus* | Short-eared Owl | TRUE |
| *Spatula clypeata* | Shoveler | TRUE |
| *Spinus spinus* | Siskin | FALSE |
| *Gallinago gallinago* | Snipe | TRUE |
| *Burhinus oedicnemus* | Stone-curlew | TRUE |
| *Saxicola rubicola* | Stonechat | TRUE |
| *Anas crecca* | Teal | TRUE |
| *Anthus trivialis* | Tree Pipit | TRUE |
| *Passer montanus* | Tree Sparrow | TRUE |
| *Aythya fuligula* | Tufted Duck | TRUE |
| *Streptopelia turtur* | Turtle Dove | TRUE |
| *Linaria flavirostris* | Twite | TRUE |
| *Rallus aquaticus* | Water Rail | TRUE |
| *Oenanthe oenanthe* | Wheatear | TRUE |
| *Saxicola rubetra* | Whinchat | TRUE |
| *Mareca penelope* | Wigeon | FALSE |
| *Poecile montanus* | Willow Tit | TRUE |
| *Phylloscopus sibilatrix* | Wood Warbler | TRUE |
| *Scolopax rusticola* | Woodcock | TRUE |
| *Lullula arborea* | Woodlark | TRUE |
| *Motacilla flava* | Yellow Wagtail | TRUE |

#### Species Habitat Associations

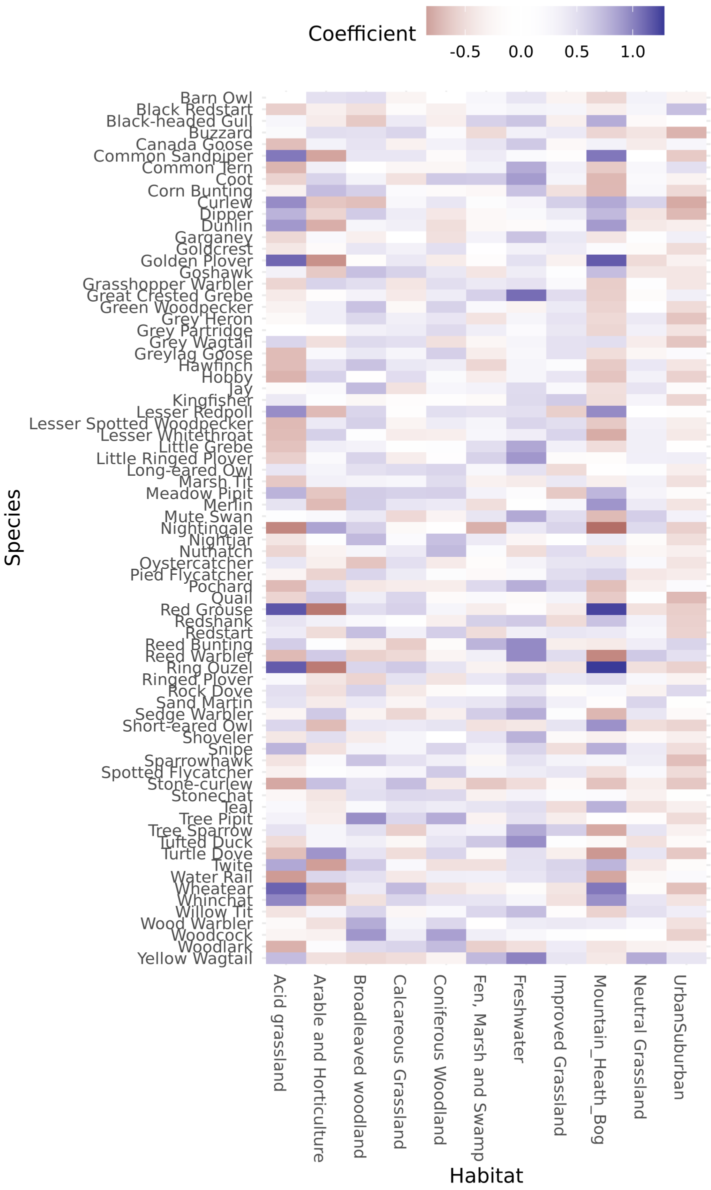

Figure 13: Fitted species-level environmental coefficients for 1970 breeding bird dataset (excluding ‘possible’ records)

#### Species Codistribution

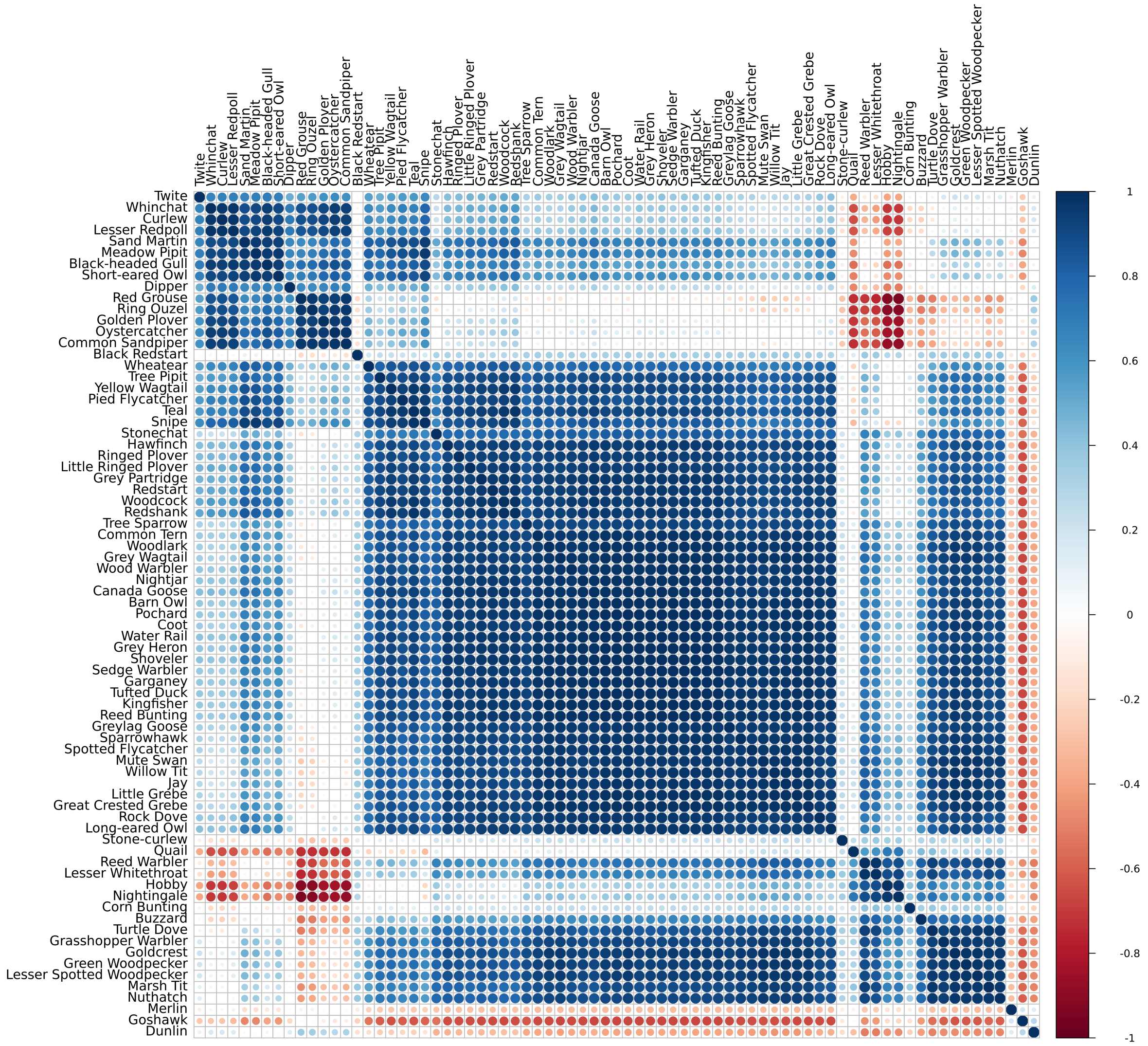

Figure 14: Fitted residual species-association matrix (correlations) for 1970 breeding bird dataset (excluding 'possible' records.)

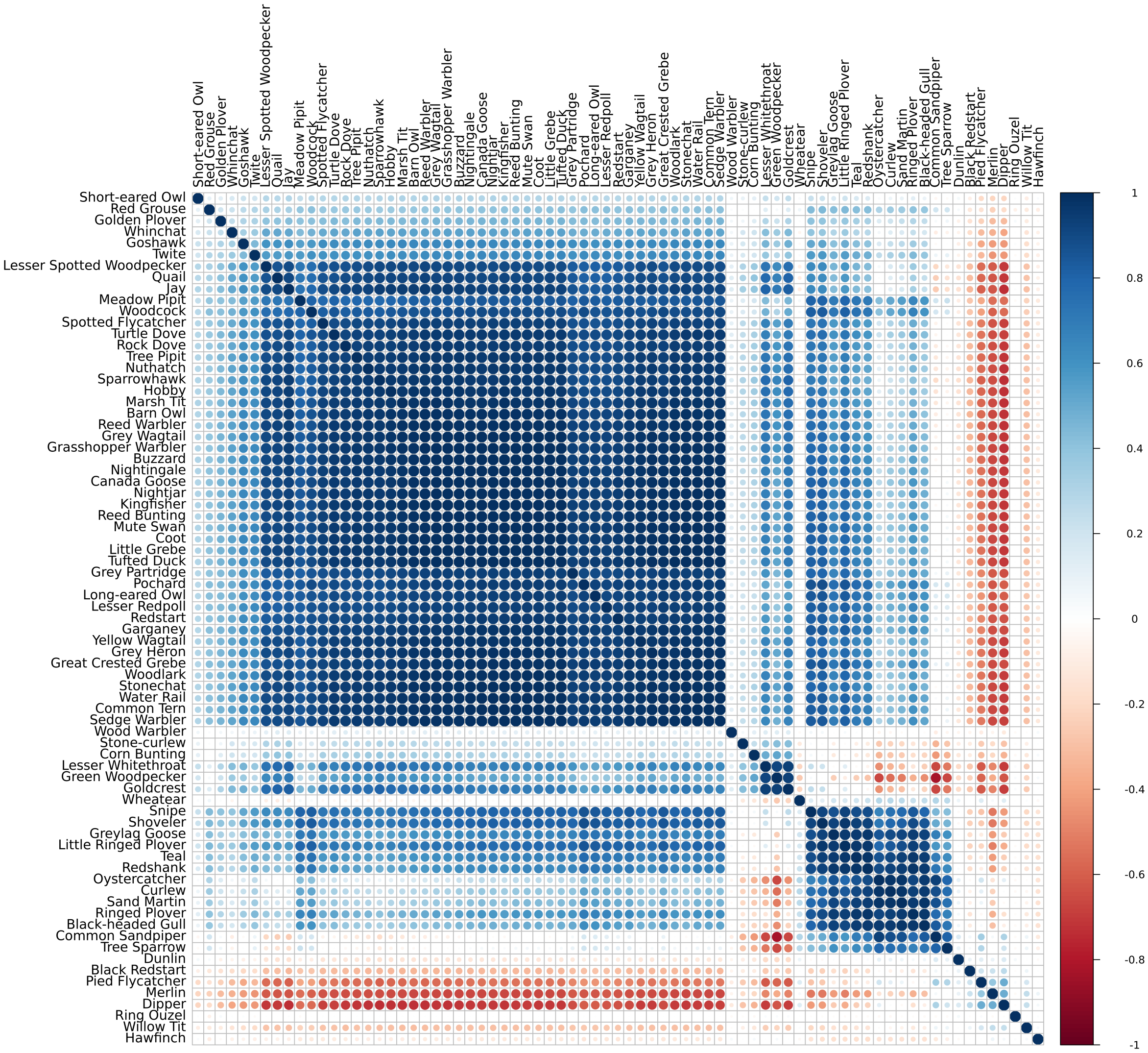

Figure 15: Fitted residual species-association matrix (correlations) for 2010 breeding bird dataset (excluding 'possible' records.)

#### MCMC Convergence

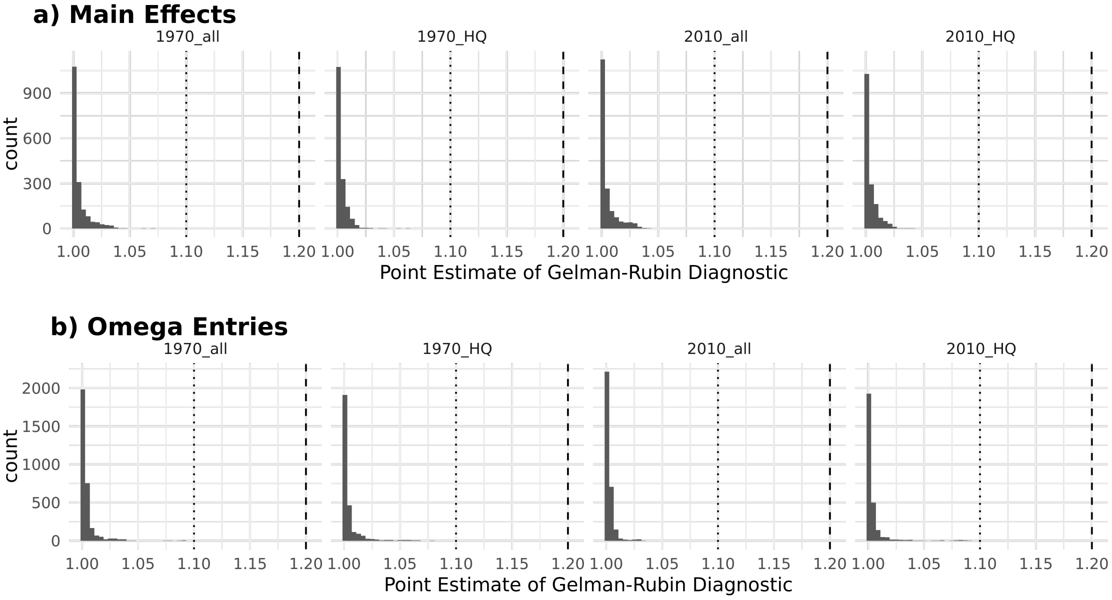

Figure 16: Distribution of Gelman-Rubin MCMC convergence diagnostic value point estimates for the bird datasets, calculated from 4 independent MCMC chains of the 'full' model. Values are faceted by year and whether all data is used, or excluding 'possible' observations (HQ). a) Main effect coefficients (i.e. environmental and spatial variables). Largest value was 1.069. b) Elements of the correlation matrix ($\Omega$). Largest value was 1.102. All were well below the standard thresholds indicating convergence is likely achieved.
